# Illusory tilt does not induce optostatic torsion

**DOI:** 10.1101/2025.07.23.666363

**Authors:** Mihret Girum, Ariel Winnick, Jorge Otero-Millan

## Abstract

Viewing tilted images that contain spatial information about gravity and verticality, such as rotated landscapes, or leaning buildings, produces an eye movement response known as optostatic torsion (OST). OST consists of a small-amplitude rotation ∼1º of the eyes around the line of sight, in the same direction of image tilt. Here we aimed to determine whether illusory perceptions of visual tilt were sufficient to produce OST. The illusory stimulus was a variation of the Café Wall illusion, applied to four “walls” to approximate the appearance of a tilted room. In a first experiment we determined the perceived magnitude of tilt of the illusory stimulus using a two-alternative forced choice (2AFC) task, measuring the amount of tilt required to cancel the perceived illusory tilt for clockwise (CW) or counterclockwise (CCW) configurations. We found an illusory tilt of +3.67±0.57° and -3.80±0.93°, respectively. Then, in a second experiment, we recorded 3 dimensions of binocular eye movements in ten healthy subjects viewing one of four possible stimuli: 1) illusory, 2) non-illusory with small tilt, 3) landscape with small tilt, and 4) landscape with large tilt. Both the landscape (±4º tilt: 0.4±0.1, p<0.05; ±30° tilt: 0.5±0.1, p<0.01;) and control stimulus (0.2°±0.1°, p<0.05) produced a significant amount of OST when comparing left tilt and right tilt configurations while the illusory stimulus (0.11°±0.07°, p=0.15) did not. This indicates a potential dissociation between our perception of tilt and the processing of tilt that drives the motor response of OST.

## INTRODUCTION

In healthy individuals, torsional eye movements, i.e. rotations around the line of sight, can be observed under several ordinary circumstances, e.g., when the eyes shift to an oblique position (Ferman et al., 1987), or when the head tilts towards the shoulders (Misslisch et al., 1994). Studying torsional eye movements has provided many clues to how the eyes and brain integrate information to create seamless visual experiences (Brodsky, 2002). Torsional eye movements have often been investigated in the context of gaze shifts, vestibulo-ocular reflexes (VOR), ocular counter-roll (OCR), and optokinetic (OKN) stimuli (Leigh and Zee, 2015). However, the relationships between human visual perception and ocular motor pathways in the context of torsional eye movements have yet to be well understood.

Several studies have observed optically induced torsion of the eyes, also called optostatic torsion (OST), when viewing static tilted stimuli, on the order of 1° (Mesker, 1953; Crone, 1975). Similar presentation of static tilt using the classic rod-and-frame test stimulus was also found to induce torsional responses, ranging from 0.10 to 0.78° (Goodenough et al., 1979). In contrast, some studies have found no torsion from viewing a static tilted line, only observing a response when the tilted line rotated slowly at 0.8-8° per second (Howard and Templeton, 1964). Other studies have found no torsion from viewing a static horizontal grating but measured torsional deviations when the gratings oscillated in the frontal plane at 0-0.6 Hz, with maximal torsion at 0.1-0.2 Hz (Kingma et al., 1997). Although there has been variability in early findings of optostatic torsion, it has been demonstrated that stationary visual stimuli can drive ocular torsion, but to a lesser degree than optokinetic visual stimuli.

To further explore these torsional responses driven by optostatic stimuli, more recent studies have investigated the properties of these stationary tilted visual stimuli that seem to induce ocular torsion. For example, photographic visual scenes with familiar spatial clues such as buildings, waterfronts, and the sky, were found to induce more ocular torsion than stimuli without such spatial clues, as measured quantitatively by 3D video oculography (VOG) (Pansell et al., 2006). This suggested that torsional responses may be greater with the addition of familiar spatial information. Interestingly, in this study, even a simple patterned rectangular image induced ocular torsion when viewed tilted at 15, 30, or 45°. Such evidence suggests that simple geometric patterns can also induce optostatic torsion.

With the understanding that visuospatial information can drive a torsional response, recent studies have begun investigating the link between ocular torsion and perceptual illusions. These experiments imply dynamic links between conscious perception and ocular torsion. For static viewing conditions, it seems likely that higher-order perceptual processes may influence the torsional eye position, but these phenomena require further investigation (Sverkersten et al. 2009).

Here we aimed to determine whether illusory perception of static tilt was sufficient to produce OST. To ask this question, we sought a visual stimulus that could produce the feeling of being in a tilted room without using any objectively tilted lines. We identified a visual stimulus from a family of static tilt illusions called Café Wall type illusions that influence the perception of tilted lines (Kitaoka et al., 2004). This particular image (Figure 1A) also resembled the non-illusory stimulus found to induce ocular torsion by Pansell et al., 2006. It is composed of alternating black and white squares and is thought to produce an illusory perception of tilt by combinations of contrast polarities (Kitaoka et al., 2004). Such illusory tilts are typical of Café Wall type illusions and have been explained by various analyses of luminance and border-locking of the alternating black and white squares (Gregory and Heard, 1979). Numerous other types of static tilt illusions and their properties (e.g. Zöllner, Fraser, and Haig illusions) have been reviewed elsewhere (Kitaoka, 2007). The illusory image chosen for this investigation induced the perception of tilted lines and had an additional spatial component that may induce the sense of being inside a room or corridor with a floor, ceiling, and walls. We wondered if this stimulus could engage the brain’s spatial representations of tilt without any objective visual, vestibular, or somatosensory stimulus to drive a tilt signal.

**Figure 1.**
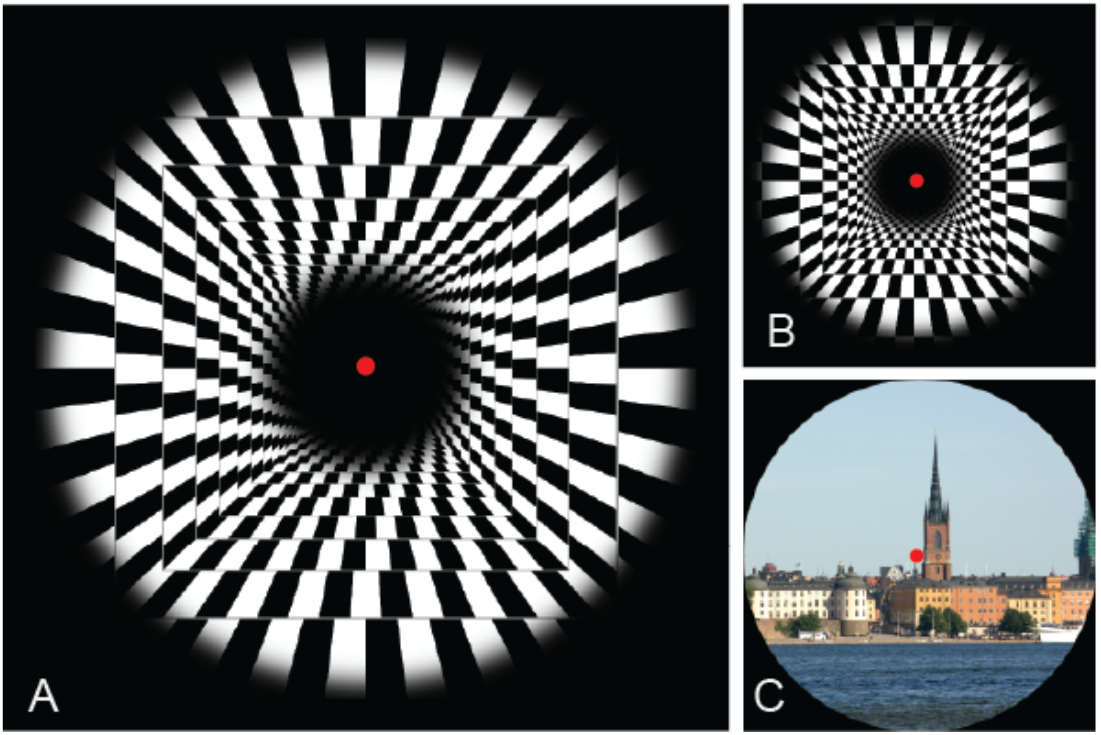
The three visual stimuli used in the experiment, shown in neural orientation (0**°**). **A)** variation of the Café Wall illusion (Kitaoka et al., 2004) inducing an illusion of rightward tilt. **B)** a symmetrical variation of A that does not induce an illusion of tilt. **C)** Natural scene stimulus with spatial cues utilized for comparison.

**Figure 2.**
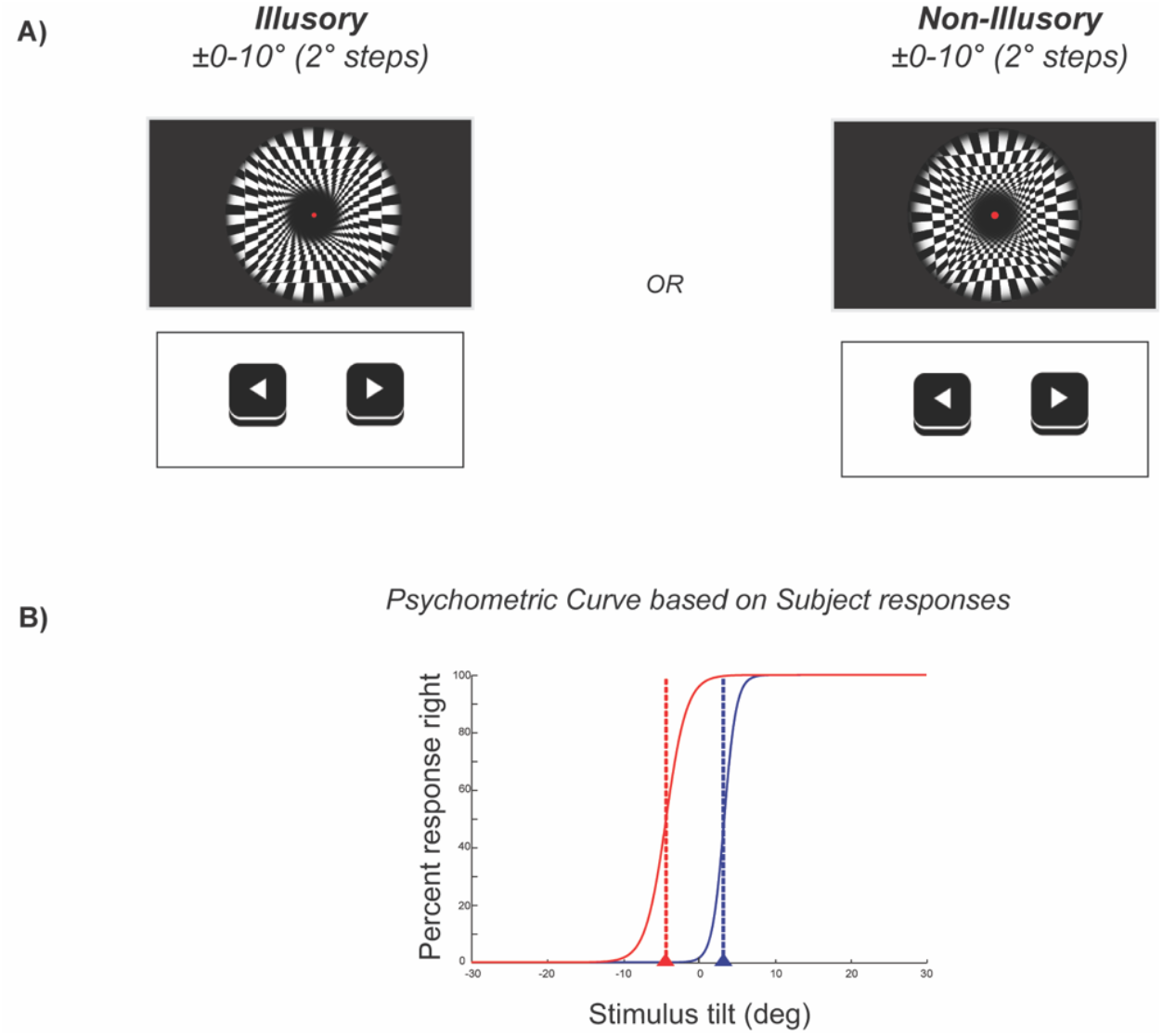
Experiment 1 Sequence: determined the amount of tilt perceived by each subject using the illusory stimulus and comparing it to the non-illusory control stimulus. **A)** Displays the experimental design and the two stimuli presented. **B)** is an example of an SVV psychometric curve generated by a single subject for the illusory stimulus.

To test the possible influence of static illusory tilt on torsional eye position, we asked whether visual illusion of tilt in the frontal plane can reliably drive optostatic torsion. We measured torsional eye position with subjects viewing the illusory tilted room in the upright position (0°); to create the illusion of both directions of tilt (CW and CCW), we used contrasting polarities.

The effects of the static illusion were quantified by comparing measured eye movement responses to both illusory images and non-illusory tilted images.

## EXPERIMENTAL PROCEDURES

### Subjects

We tested 10 healthy subjects (28.2 ± 6.1 years, 4 males) in experiment 1, and 10 healthy subjects (24.7 ± 1.6 years, 2 males) in experiment 2. All participants reported normal vision (>20/100 uncorrected), no history of ophthalmologic or vestibular disease and provided written informed consent prior to participation. The experimental protocol was approved by the Committee for Protection of Human Subjects (CPHS/IRB) of the University of California, Berkeley.

### Set-Up

Participants were seated in front of a 4K OLED screen (3840 x 2160 pixels, 144.4 x 82.2 cm, 250 Hz refresh rate, LG OLED65CXPUA), with a 90cm eye to screen distance and the head stabilized using an adjustable headrest. The experimental paradigm used to present visual stimuli was programmed in MATLAB Version R2021a (MathWorks, Natick, MA) using Psychtoolbox (Kleiner et al., 2007). All stimuli were displayed against a black background in a dark room.

### Stimuli

Participants were presented with three types of stimuli. The illusory stimulus was a variation of the Café Wall illusion (Kitaoka et al. 2004) extended to four walls to provide spatial cues and produce the appearance of a tilted room, both right and leftwards configurations were presented. This illusion utilizes offset black and white tiles separated by “mortar” lines to create an illusion of a slanted line. The lines separating the tiles (“mortar” lines) were 50% gray between the white and black tiles. The non-illusory stimulus consisted of a similar pattern of offset black and white tiles, without the “mortar” lines. The last stimulus was a landscape image of a city scene with spatial cues. The non-illusory and landscape stimuli were presented in CW and CCW tilt orientations, the magnitude of tilt was determined by Experiment 1.

### Experiment 1

To assess perceived tilt of the illusory stimuli, participants were presented with either an illusory or non-illusory stimuli (Figure 1). A brief introduction and demonstration were provided to familiarize participants with the psychophysical task and allow participants to practice reporting their perceived orientations of the images. Each paradigm presented a central red fixation dot (5mm, 0.32° visual angle) coincident with stimuli presentation. A sequence of illusory or non-illusory stimuli were then presented in a random order of angles ranging from 0-10^°^ CW or CCW in ±2° increments. Participants were instructed to report whether the images looked tilted right or left by pressing corresponding arrow keys on the keyboard, stimuli remained visible until a selection was made. Sequences of illusory (52 cm, 32°) and non-illusory (81 cm, 48°) stimuli were presented separately, into two sessions (a single illusory and non-illusory session) of 100 trials each.

### Experiment 2

To measure the torsional responses to the stimuli, subjects were equally presented with eight stimuli in a random order for a total of 64 trials. The stimuli consisted of: natural scene images presented in both small (±4°) and large tilt (±30°) configurations, non-illusory images presented in small tilt configurations (±4°), and the illusory images presented in both right and left tilt configurations (Figure 3). Each trial was initiated with 5 seconds of central fixation with a red fixation dot (5mm, 0.32° visual angle), the stimulus was subsequently superimposed for 15 seconds while the subject maintained central fixation with awareness of the full stimulus.

**Figure 3.**
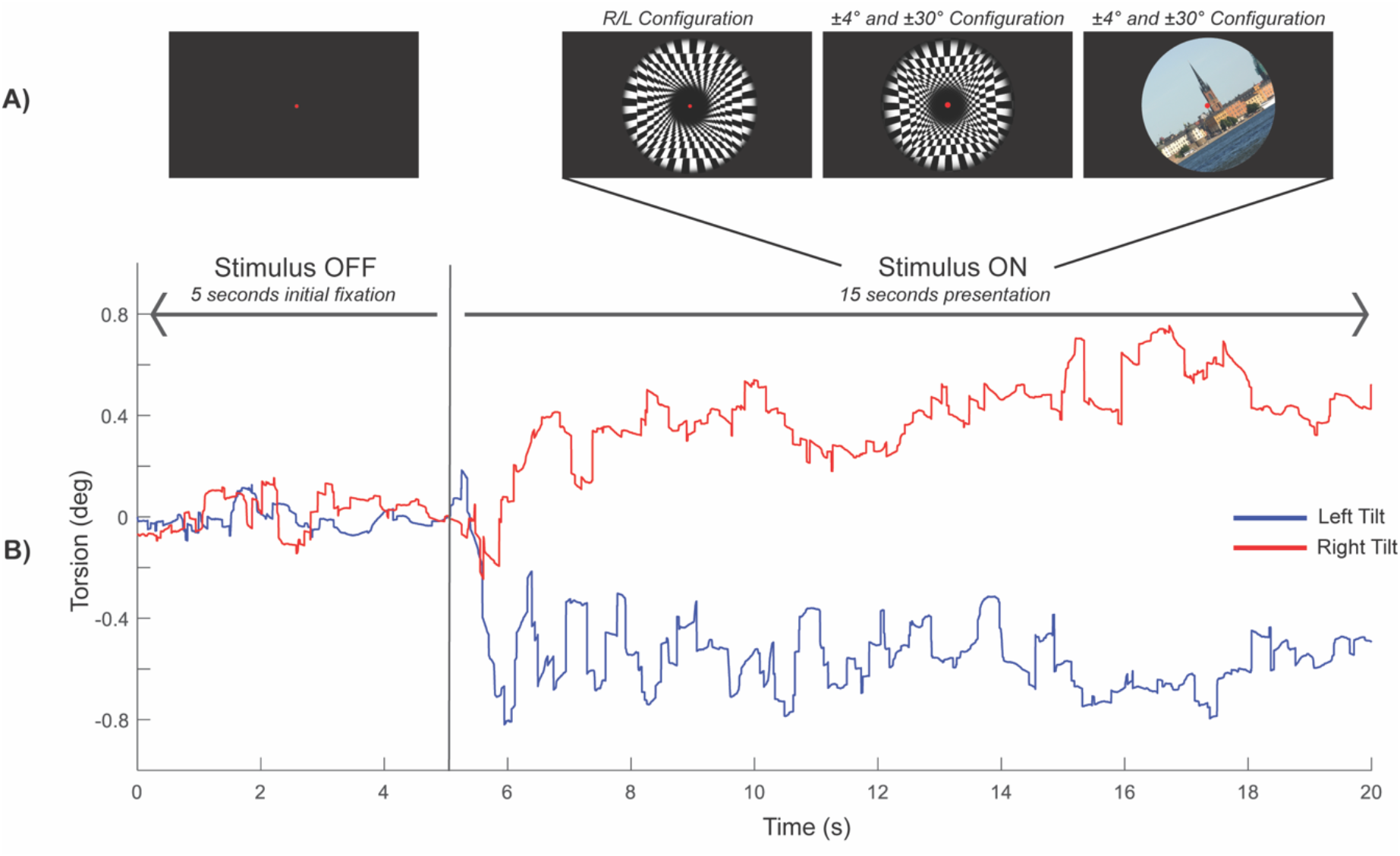
Experiment 2 measured torsion induced by attending to the illusory, non-illusory and natural scene stimuli while maintaining fixation. **A)** Displays the experimental sequence, configurations of the stimuli and duration each component was presented to the subject. **B)** E xample of raw torsional eye movement recording from a subject during the experiment.

Throughout the trials eye movements were recorded and processed to determine the presence of ocular torsion during stimulus presentation.

### Eye Movements Recordings and Analyses

Eye movements were recorded (250Hz) binocularly using an infrared camera (FLIR Grasshopper3 USB3), placed 22.7cm away from subjects, and using the custom software that tracks the pupil and the iris to measure eye movements in three dimensions. To limit interference with recordings, subjects were asked to remove any eyeglasses. Participants whose uncorrected visual acuity did not allow resolution of the images on screen were excluded from the study.

Torsional movements were measured by amount of shift from each subject’s reference iris template established during calibration (Otero-Millan et al. 2015).

### Statistics

Data from the perceptual experiment was averaged based on each participant’s SVV, magnitude of tilt required to neutralize the illusion, for both CW and CCW orientations of the stimulus. A t-test analysis was then used to determine the statistical significance. The data from the torsional experiment was averaged for each stimulus type (landscape ±4° and ±30°, illusory ±4°, and non-illusory R/L) combining responses to positive tilts and inverted responses to negative tilts. We then used another t-test analysis for each stimulus type to determine the statistical significance of the measured torsion response.

## RESULTS

### Perceptual Experiment

The magnitude of perceived tilt was determined by fitting a psychometric curve to the subjects’ responses to determine at which tilt angle the stimulus was perceived as upright (chance rate of CW or CCW responses (Figure 1B)). This value indicated the amount of actual image tilt required to cancel the perceived illusory tilt of the stimulus. In addition to the illusory stimulus, control stimuli were also presented to the subjects.

Figure 4 displays a summary of the average amount of tilt at which the illusory stimuli were perceived to be neutral or upright. The CW and CCW configurations of the illusory stimuli were perceived to be upright with -3.80±0.93° and 3.67±0.57° tilt, respectively. This indicates that the illusory stimuli generated a perceived tilt of approximately 4° on average by subjects. We used an approximation of this value of 4° to decide how much to tilt the non-illusory and landscape stimuli in Experiment 2 as control stimuli. The non-illusory stimuli, when presented at CW and CCW orientation, were perceived as vertical with -1.62±1.05°(p=0.158) and 0.94±1.09° (p=0.411), respectively.

**Figure 4.**
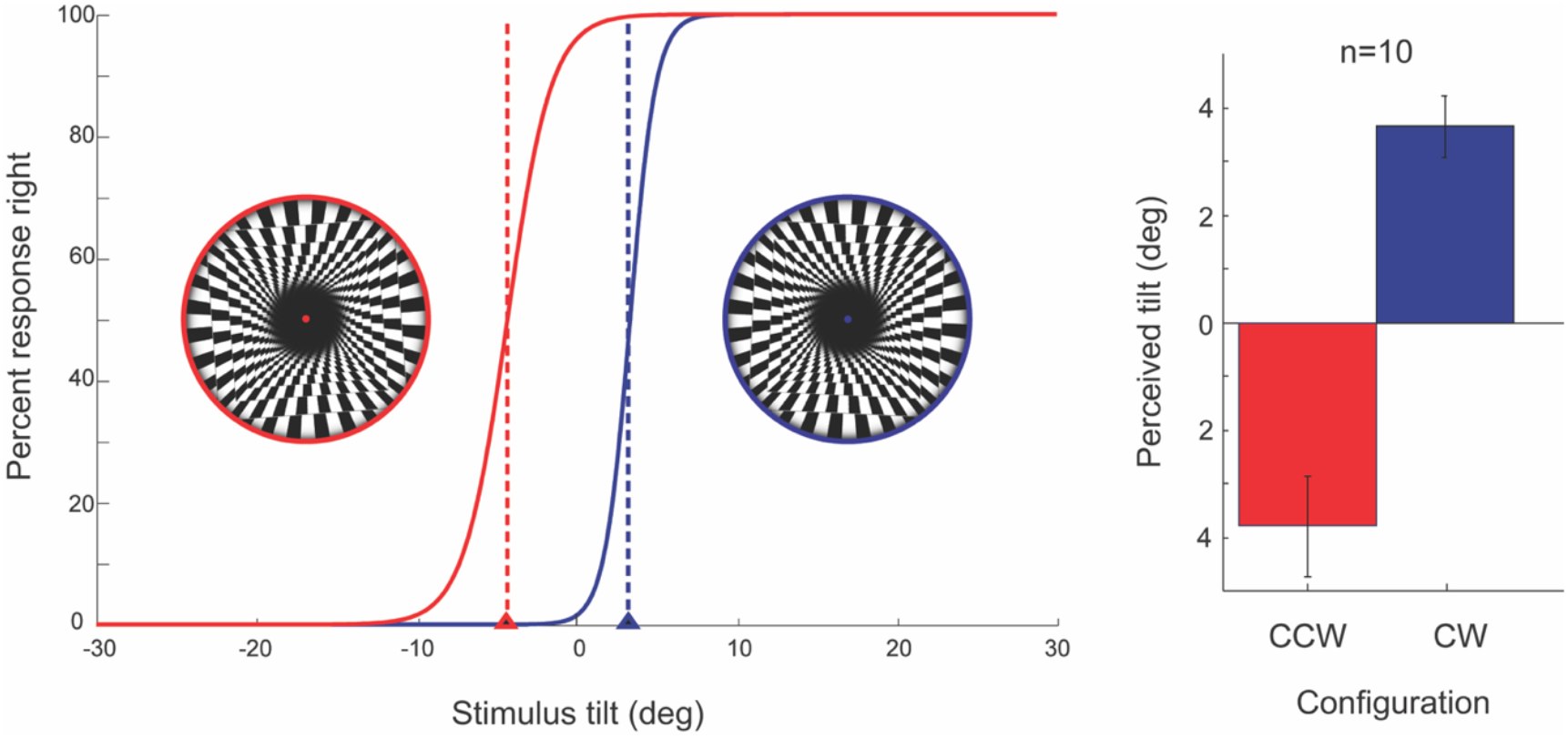
Average mean effect of the illusory stimuli (n = 10) from Experiment 1. The red denotes the CCW or left orientation of the illusion and the blue indicates the CW or right orientation of the illusory stimuli.

### Torsion Measurement

#### Effect of Illusory Stimuli

We found that all stimuli, apart from the illusory stimulus, produced a significant optostatic torsional response (OST) (Figure 5,6). The illusory stimulus did not induce a statistically significant amount of OST (0.11°±0.07°, p=0.151).

**Figure 5.**
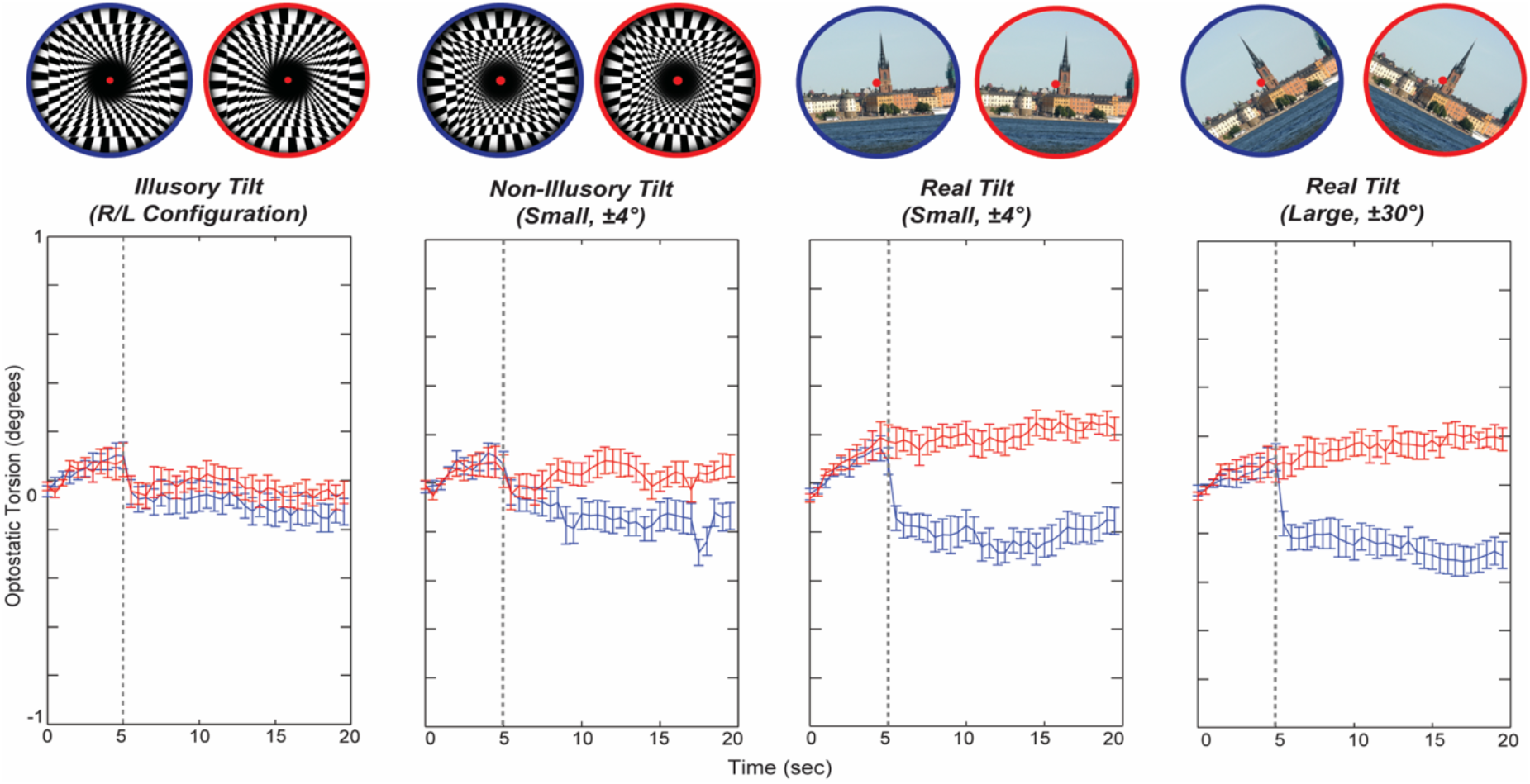
Average torsion responses for each stimulus presented (n=10). Red and blue denote stimuli oriented to the right and left, respectively. Trials consisted of an initial 5 second fixation period, grey dotted line indicates the end of this period, followed by 15 seconds of stimulus presentation for a total of 20 seconds per trial. Illusory images were presented in two configurations (right or left illusory tilt), the non-illusory control and Real Small Tilt images were titled ±4^°^, Real Large Tilt images were tilted ±30^°^. All subjects and trials of the same stimuli were averaged. Note that subjects present some initial drift and some sudden change in torsion associated with the stimulus onset regardless of the stimulus presented.

**Figure 6.**
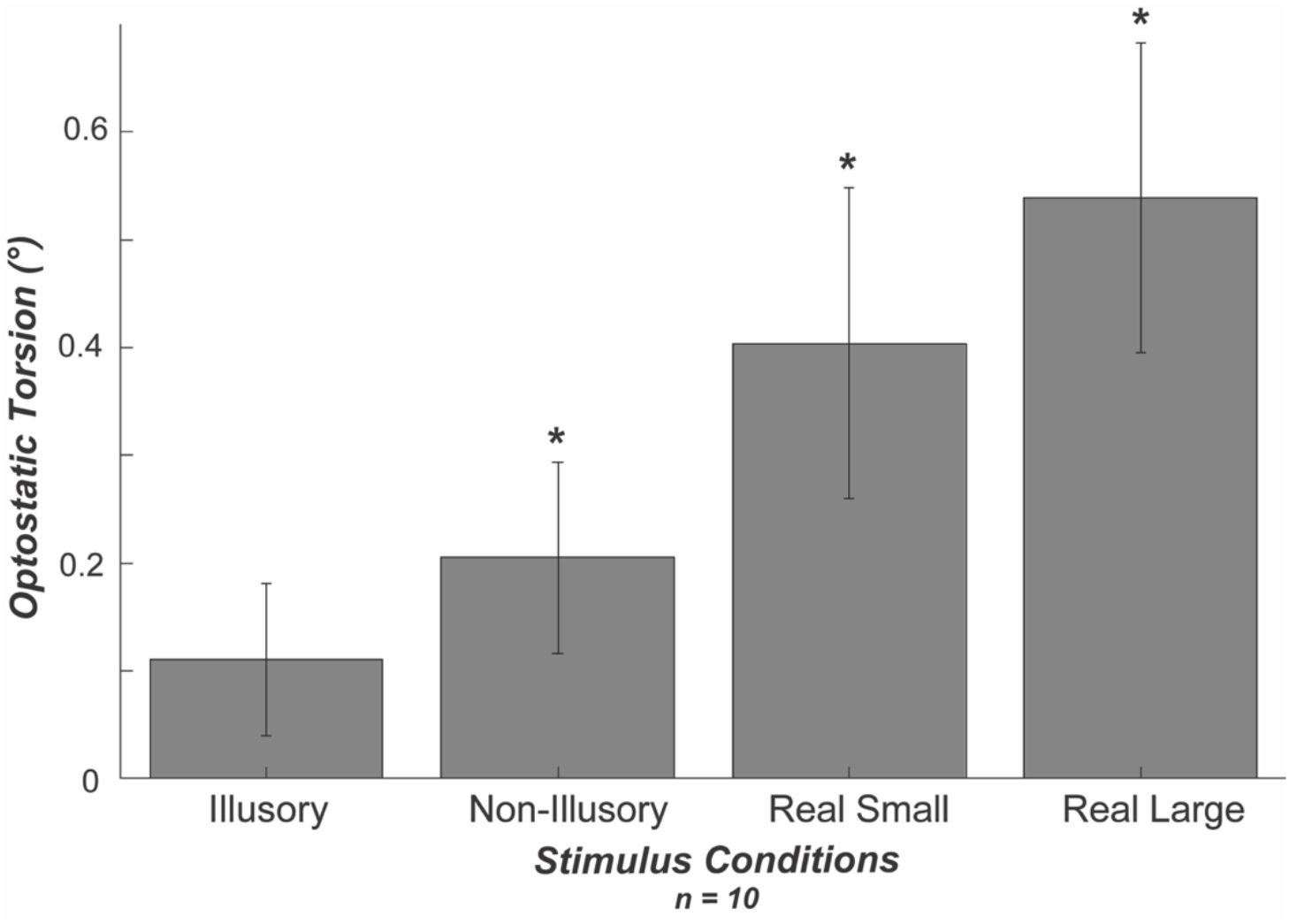
Average OST for each stimuli condition presented (n = 10) from Experiment 2. Asterisks indicate the conditions that produced a significant OST response.

#### Effect of Landscape Visual Stimuli

Comparison of the average OST responses during stimulus presentation, shows that the tilted landscape stimulus (4°: 0.43°±0.11°, p<0.01; ±30°: 0.40°±0.14°, p=0.021) was able to induce a larger OST response than the illusory stimulus (0.11°±0.07°, p=0.151).

#### Effect of Tilt Magnitude

The landscape images were presented at two magnitudes of tilt, ±4° and ±30°. We found no difference caused by the magnitude of landscape image tilt, both ±4° (0.43°±0.11°, p<0.01) and ±30° (0.40°±0.14°, p=0.021) orientations produced similar magnitudes of compensatory OST.

#### Effect of Control Non-Illusory Stimuli

To determine whether the abstract pattern had any influence on the OST response, a similar black-and-white tiled pattern without the illusory tilt effect was presented to participants. This stimulus was presented at ±4°. This stimulus did on average produced a significant compensatory OST response (0.21°±0.09°, p<0.01). However, it did not statistically differ from the minimal OST response induced by the illusory stimulus (p=0.4).

## DISCUSSION

In this study, our objective was to determine whether viewing an illusory tilted stimulus could generate an optostatic torsion (OST) response. We used a variation of the Café Wall illusion (Kitaoka et al. 2004) approximating the 4 walls of a virtual room (walls, floor, and ceiling) to provide added spatial cues. We found that healthy participants did not exhibit robust OST responses while viewing illusory stimuli, but OST was reproduceable with natural and abstract stimuli objectively tilted in the frontal/roll plane.

Optostatic torsion (OST) responses have been elicited in observers using variations of tilted static stimuli (Pansell et al., 2006, Sverkersten et al., 2009). Images of tilted landscapes, or even abstract shapes, seem to provide the brain with enough visual information to generate the perception of a tilted world. When the head is stabilized in an upright position, vestibular and proprioceptive inputs register an upright world, in which earth-horizontal and gravitational vertical correspond. Our visual experience of horizontal and vertical meridians are expected to align with other visual, vestibular, and proprioceptive inputs, as these are believed to contribute to a common internal reference frame (Angelaki and Cullen 2008). Images that indicate tilt in the fontal/roll plane thus produce sensory conflict, because vestibular and proprioceptive afferents indicate upright orientation (0°). In such cases, an internal error signal – representing a difference between visual, vestibular, and proprioceptive input – appears to be sufficient to generate OST. With a tilted image representing a false upright orientation to the visual system, the head/body is perceived as tilted in the opposite direction, just as the otolith organs would register if the body were tilted but the visual percept veridical with respect to the world. In correspondence with this internal bias, the OST is a compensatory response, a rotation about the roll-axis in the same direction of the image tilt.

In our study, participants were presented with two types of experimental stimuli: landscape and abstract. Landscape scenes provided gravity cues that stimulated compensatory torsional responses in the direction of image tilt. These torsional eye movements are likely triggered by the outcome of weighted visual, proprioceptive, and vestibular information (Kheradmand and Winnick, 2017; Ozturk et al. 2021). Within the vestibular system, perceived orientation of self, relative to the environment (i.e., with respect to gravity), is mediated by the otoliths and is a major driver of ocular torsion (Groen, Howard & Cheung, 1999). The magnitude of these compensatory responses is only a small percentage of the actual tilt presented to subjects, indicating that these torsional responses are not intended to re-align retinal coordinates but, rather provide some perceptual benefit that has yet to be well understood (Pansell et al, 2006).

The forced-choice experiment indicated that upright illusory stimuli was on average perceived to have 4° tilt (CW/CCW). Although this tilt was less than utilized in torsion response experiments by Pansell et al., 2006, we did find an OST response when the same extent of tilt was presented with the landscape stimuli. This response was limited and did not provide full compensation.

While it has been postulated that compensatory torsional responses are influenced by the brain’s preference to analyze vertical images without tilt, the limited torsional response found with image tilt suggests otherwise and further presents the limitations of ocular torsional movements (Pansell et al., 2006).

While every tilted (non-illusory) stimulus that we presented produced significant OST responses, the illusory stimulus did not. This may indicate a decoupling between perception and action as noted in many other studies (Betts & Curthoys, 1998; Dakin et al., 2020). Here we observe that a stimulus can produce a very clear and significant perceptual tilt while not inducing a corresponding motor response (OST). This decoupling could correspond with different neural pathways responsible for the perception of tilt of the external world and the pathways integrating visual and vestibular information to drive our motor reflexes, including OST. Similar decoupling has been observed between torsion and perception of upright, for example for prolonged tilts when perception changes much more dramatically than torsion during the tilt (Otero-Millan & Kheradmand, 2016). Alternatively, this variance in response may be attributed to the prominent differences of the two stimuli categories. The visual system is primed to optimally respond to natural scenes, these stimuli are often low spatial frequency requiring less effort to decode (Powell et al., 2006). Visual stimuli that deviate from typical natural scene stimuli have been found to be processed less efficiently (Parraga et al. 2000). The Café Wall illusion utilizes offset black and white tiles separated by “mortar” lines to create an illusion of a slanted line. The use of repeated tiles across the stimulus presents an image with higher spatial frequency, that can cause visual discomfort or stress, lending to limitations in visual processing (Barten, 1999; Campbell & Robson, 1968). Furthermore, illusions indicate conflict between perceptual and sensory information; inefficient decoding of an illusory stimulus may contribute to cortical sensory re-weighting processes that downregulates visual information. If so, this would increase emphasis on vestibular afferent input which would trigger minimal ocular motor response as a result of head-stabilized subject position.

In our study, we did not find a significant difference in OST response between small (±4°) and large tilt (±30°) configurations, appearing to contradict the previously proposed increases in OST with increased image tilt (Pansell et al., 2006). However, in this study, larger angles of tilt were tested (15, 30, and 45°) in comparison to the small 4° utilized in our experimental paradigm. We also do see a trend of increased OST with the larger tilt that is consistent with their results, potentially suggesting that due to the small magnitude, our findings may have been negligible in our sample. However, it does suggest that OST may initially increase fast as a function of image tilt angle and then vary little with larger tilt angles. It is also possible that in instances of conflict between visual and vestibular information, cortical processes will reweight sensory information to align with reliable signals (Ozturk et al., 2021; DiGirolamo et al., 2001; Dakin et al., 2020; ter Host et al., 2015). Increasing magnitude of natural scene tilt may influence a similar process causing downregulation of visual information integration due to disagreement with upright head-stabilized vestibular input, thus maintaining a consistent response with varying image presentations.

In summary, we have found an additional dissociation between perception and motor action. The way in which our brain extracts a sense of tilt from a visual image affects differently our perception of tilt and our ocular motor reflexes. This may provide insights into the mapping of sensorimotor pathways as well as in developing diagnostics tests that differentially probe motor and perceptual deficits.

## ACKNOWLEDGEMENTS

This project was funded by NEI Training grant 5T32EY007043-43 and NEI R00EY027846.

